# Two modes of mid-frontal theta suggest a role in conflict and error processing

**DOI:** 10.1101/2022.03.25.485421

**Authors:** Vignesh Muralidharan, Adam R Aron, Michael X Cohen, Robert Schmidt

## Abstract

Mid-frontal theta increases during scenarios when conflicts are successfully resolved. Often considered as a generic signal of cognitive control, its temporal nature has hardly been investigated. Using advanced spatiotemporal techniques, we uncover that mid-frontal theta occurs as a transient oscillation or “event” at single trials with their timing reflecting computationally distinct modes. Single-trial analyses of electrophysiological data from participants performing the Flanker (N = 28) and Simon task (N = 18) were used to probe the relationship between theta and metrics of response conflict. We specifically investigated “partial errors”, in which a small burst of muscle activity in the incorrect response effector occurred, quickly followed by a correction. We found that transient theta events in single trials could be categorized into two distinct theta modes based on their relative timing to different task events. Theta events from the first mode occurred briefly after the task stimulus and might reflect conflict-related processing of the stimulus. In contrast, theta events from the second mode were more likely to occur around the time partial errors were committed, suggesting they were elicited by a potential upcoming error. Importantly, in trials in which a full error was committed, this “error-related theta” occurred too late with respect to the onset of the erroneous muscle response, supporting a role of theta also in error correction. We conclude that different modes of transient mid-frontal theta can be adopted in single trials not only to process stimulus conflict, but also to correct erroneous responses.

## INTRODUCTION

Resolving response conflict helps us select appropriate actions among several competing ones. The neural mechanisms associated with conflict and error processing have been linked to increases in theta-band activity (4-8Hz) in mid-frontal brain regions (Cavanagh & Frank, 2014; Cohen & Cavanagh, 2011; Cohen & Donner, 2013; Cohen & van Gaal, 2014; Nigbur, Cohen, Ridderinkhof, & Stürmer, 2012) and to the coherence in the theta-band between mid-frontal cortex and basal ganglia, specifically implicating the hyperdirect pathway (Aron, Herz, Brown, Forstmann, & Zaghloul, 2016; Zavala et al., 2016). Theories on mid-frontal control processes have related theta activity increases to the need for control, which arises in situations requiring the resolution of a conflict at the level of stimulus, response, or even rewards (Cavanagh, Figueroa, Cohen, & Frank, 2012; Cavanagh et al., 2011; Cavanagh & Frank, 2014; Frank, Woroch, & Curran, 2005). Mechanistically, this has been attributed to theta’s ability to modulate reaction times by allowing extra time for evidence accumulation by increasing decision thresholds (Cavanagh et al., 2012, 2011; Frank et al., 2005). However, while theta increases are seen in trials in which conflicts were resolved successfully, they also occur in trials after an error was committed, suggesting multiple roles of theta with potentially different neural sources (Swick & Turken, 2002). With many of these findings looking at average theta power across conditions and their relationship to underlying metrics of conflict, the nature of theta at the single-trial level, specifically the time at which it occurs and the temporal relationship to conflict measures is still not known. Therefore, we investigated how the temporal dynamics of theta within a trial relate to an impending error and to the processes that allow the correct response to get elicited.

In response-conflict paradigms, “partial error” trials provide additional insights into the amount of response conflict experienced by the research participants in single trials. In partial error trials, the participants started to initiate an incorrect response (measured through electromyogram or force grips), but then quickly corrected the erroneous action to make a correct response (Cohen & van Gaal, 2014; Rochet, Spieser, Casini, Hasbroucq, & Burle, 2014; Vidal, Burle, & Hasbroucq, 2019). In these partial error trials, mid-frontal theta power has been linked to the changes in level of conflict experienced, indexed by the amount of time it takes to correct the error (Cohen & van Gaal, 2014; Cohen, 2014), although a clear temporal picture of when each event happens within a trial remains unclear.

In addition to the improved characterization of response conflict in single trials, advances in the analysis of oscillatory signals have provided a more accurate picture of the temporal dynamics in single trials. In contrast to oscillations that are ongoing and slowly modulated in their amplitude, several recent studies have revealed that average increases in oscillatory power in a particular frequency band can be due to transient burst-like oscillations in single trials (Feingold, Gibson, DePasquale, & Graybiel, 2015; Little, Bonaiuto, Barnes, & Bestmann, 2019; Torrecillos et al., 2018). For instance, transient beta oscillations, also referred to as beta bursts, in motor and frontal cortical areas are more tightly linked to behavior compared to trial-averaged power increases (Little et al., 2019; Muralidharan, Aron, & Schmidt, 2021; Muralidharan & Aron, 2021; Soh, Hynd, Rangel, & Wessel, 2021). More generally, oscillatory bursts have been seen in other frequency bands, for example, both gamma and beta bursts have shown to predict behavioral variability during working memory tasks (Lundqvist et al., 2016). Thus, it is intriguing to ask the question whether theta oscillations also occur as transient burst-like events during response conflict scenarios. Here we used this approach to examine the relation between theta oscillations and single-trial measures of response conflict provided by partial errors.

In this study we analyzed EEG data from two well-known paradigms known to elicit response conflict: the Flanker task and the Simon task. We first found that average power changes in conflict-related mid-frontal theta can be decomposed into single-trial theta “events”. We then investigated the relationship between these theta events and muscle-level metrics of conflict, i.e. partial errors at a single trial level. We found that theta events could be divided into two modes when a partial error occurred. While one mode of theta might be linked to the stimulus conflict, in line with a role of theta in conflict resolution (Cohen & Cavanagh, 2011; Cohen & Donner, 2013; Cohen & van Gaal, 2014; Cohen & Ridderinkhof, 2013), the second mode was strongly linked to partial errors, pointing to a possible role of theta not only in the detection or monitoring of errors, but also in their correction. Overall, the results provide evidence that transient, mid-frontal theta events relate to cognitive and motor processes of conflict induced error and its correction.

## METHODS

### Participants

There were in total forty-six participants in the study, with 28 performing the Flanker task (20 females; Age range: 20-27) and 18 performing the Simon task (11 females; Age range: 20-25). Subjects were recruited from the University of Amsterdam community, and participated in these studies in exchange for course credit or 14 Euros. Each study was approved by the local ethics committee and subjects signed an informed consent document. Subjects had normal or corrected-to-normal vision and were self-reported free of neurological disorders and history of physical head trauma. Four participants from the Flanker task and three participants from the Simon task were excluded due to extremely noisy muscle activity which led to sub-optimal detection of trials with partial errors and their onset timings.

### Tasks and Data

We analyzed behavior and neural data from two different tasks which are known to induce response conflict: the Flanker task and the Simon task. Non-overlapping results from these data have been published previously (Cohen & Ridderinkhof, 2013; Cohen, 2015). The Flanker task was a modified version of the standard task, where participants responded to a central target which was either the letter T or I with their right or left thumb (Appelbaum, Smith, Boehler, Chen, & Woldorff, 2011). They were surrounded by distractor stimuli, two on either side of the central target such that there were three stimulus conditions. First was that the distractors were the same as the central target, or bilaterally congruent. Secondly, the distractor could be either left or right incongruent or partially incongruent, i.e., flanked by distractors different from the central target on either the right or left side respectively. Finally, the flanked distractors were all different from the central target or bilaterally incongruent. Conflict in this task is created due to the presence of the different surrounding stimuli. The stimulus was presented for 150ms, and the inter-stimulus interval was 1300-1700 ms. There were 1500 trials in total (40% bilaterally congruent, 40% partially incongruent and 20% bilaterally incongruent).

The Simon task was a simple visual association with a colored circle mapping onto either a right or a left response (Cohen & Ridderinkhof, 2013). The colored circle stimulus was presented on either the right or left side of a fixation cross, thus leading to visuo-spatial conflict. Conflict and stimulus repetition effects were balanced by using stimuli of four different colors.

In this task, conflict occurs when the correct response is opposite to the side the stimulus is presented in, for instance a blue-colored circle presented on the left side of the screen which requires a right-hand response, also called the Incongruent trials. If present on the same side, they are the Congruent trials. Stimuli were presented on the screen for 200ms with 1000ms to respond and a varying inter-trial interval of 700-1200ms. There were 1000 trials in total (500 Congruent and 500 Incongruent trials).

### Electroencephalography and Electromyography

EEG/EMG acquisition and analysis procedures were the same across both studies. EEG data were acquired at 512 Hz from 64 channels placed according to the international 10–20 system, and from both earlobes, using Biosemi equipment. Electromyographic (EMG) recordings were taken from the flexor pollicis brevis muscle of each thumb using a pair of surface electrodes, placed on a subject-by-subject basis approximately 5 mm apart on the thenar eminence.

### Data Analysis

All analyses were performed using EEGLAB13 and MATLAB R2016b.

#### Behavior and Partial Errors

The reaction time was estimated for the button press responses; hereafter referred as response time (RT). Response times below 200ms and above 1000ms were considered as outliers. A trial was classified as a correct or an error trial based on the response outcome, which we used to compute the accuracy for a given stimulus condition. We then identified the partial error trials using an algorithmic approach in line with a previous study (Cohen & van Gaal, 2014). We took the derivative of the EMG traces from both hands, Z-transformed across the entire trial period and then rectified. This controlled for several things, including differences in impedance and signal strength across subjects and hands. A trial was then classified as a partial error if the differentiated and z-normalized EMG trace of the incorrect hand in that trial exceeded two standard deviations within the time of the stimulus onset and the response time. Additionally, the peak of the partial error had to be more than twice the maximum of the baseline activity seen 300ms prior to the stimulus onset. Using this approach, we could reduce the chance for detecting partial errors which may have occurred due to noisy EMG. We will refer to these trials as partial error trials. We then estimated the EMG onsets of the partial errors and correct response, by first starting at the time of peak of both the EMG traces and back-tracking till the EMG trace for the corresponding hand fell below 20% of the peak for five consecutive milliseconds. In cases where the peak did not fall below 20% till the onset of the stimulus, we used 30% as the threshold to determine EMG onsets. Given that the number of trials with partial errors within a participant is low (~20%), for comparison we picked the correct trials which were matched for both the response times and stimulus type (for instance the bilaterally incongruent, partially incongruent and bilaterally congruent in the Flanker scenario). To do so for each detected partial error trial, we picked a correct trial which had the closest response time to it and had the same stimulus type. We will refer to these trials as the full-correct matched trials. Finally, there were trials where participants gave the erroneous response, which we refer to as the full-error trials.

#### EEG

Offline, continuous EEG data were high-pass filtered at 0.5 Hz and epoched from −1.5 to +2 s surrounding stimulus onset of each trial. All trials were visually inspected and those containing facial EMG or other artifacts not related to blinks were manually removed. Independent components analysis was computed using EEGLAB software (Delorme and Makeig, 2004), and components containing blink/oculo-motor artifacts or other artifacts that could be clearly distinguished from brain-driven EEG signals were subtracted from the data.

#### Spatial Filtering using GED: Mid-frontal source

In both task datasets, we performed a guided source separation analysis, i.e., generalized eigen decomposition, GED (Blankertz, Tomioka, Lemm, Kawanabe, & Muller, 2008; Cohen, 2017, 2022; Parra & Sajda, 2003), to obtain a mid-frontal spatial filter for extracting the conflict-related theta activity. GED allows for a more guided approach to spatial filtering by allowing us to estimate a spatial filter based on previous hypotheses regarding the spatial topography, frequency band and time-period from the data. Here, we were interested in mid-frontal theta activity (4-8Hz) around the time where conflict related processing would occur (i.e., in relation to the stimulus onset, 200ms to 100ms after the mean RTs in the high conflict condition). Thus, by pooling all correct trials and filtering the data in theta frequency (4-8Hz), we applied GED with respect to the broadband data, i.e., GED was performed between the theta filtered covariance matrix and the broadband covariance matrix (using the *eig* function in MATLAB). This provided us with spatial filter weights, aka GED components, which maximized the signal to noise ratio in theta frequency band. These weights were then forward-projected onto the theta-filtered covariance matrix to obtain spatial activation maps to visualize each component’s scalp distribution. We then selected the filter having a mid-frontal spatial distribution. In scenarios where more than one mid-frontal filter was obtained, we picked the one which explained the maximal variance in the data, as estimated using the generalized eigenvalues. Finally, the epoched EEG data were projected onto the selected mid-frontal spatial filter for time-frequency as well as theta event analyses.

#### Time-Frequency analyses

Time-frequency analysis was done in EEGLAB13 using the *newtimef* function. For the event-related spectral perturbations (ERSPs), the time-frequency decomposition was done using Morlet wavelets with frequencies ranging from 4-30Hz. We started with 3 cycles at the lowest frequency, linearly increasing as the frequencies increased such that it was 11.25 cycles at the highest frequency. The ERSPs were baselined to a period prior to the stimulus (−500 to −100ms in both task datasets). From the ERSPs of the condition potentially capable of evoking maximal response, i.e. the bilaterally incongruent trials in the Flanker and Incongruent trials in the Simon paradigm, we estimated at an individual level the peak-theta frequency within the stimulus to the corresponding condition’s mean RT. Our rationale for this approach was to select a participant-specific theta which would best reflect the underlying conflict-related processing. We used the peak-theta frequency value to extract single-trial theta events for each participant.

#### Extracting Single-trial Theta Events

From the GED component time series, we extracted single-trial theta events by first applying a temporal filter. The filter was a frequency-domain gaussian with a peak at the individual’s peak-theta frequency and a full-width half maximum of 5 Hz. We found that this was a good tradeoff between obtaining frequency specificity and temporal precision. The complex analytic time-series thus obtained was subjected to a Hilbert transform to compute its amplitude/envelope. From the theta amplitudes in a baseline period (−500ms to −100ms prior to the stimulus onset), we derived threshold values to ascertain which periods of time could be classified as a theta event. Firstly, we used an event definition threshold (median + 1.75SD). Any theta amplitude above this threshold was considered a theta event. Next, once all the theta events in a trial were evaluated, we lowered the threshold value in order to estimate both the timings and durations of the events identified using the higher threshold, i.e., an event length threshold (median +1.0SD). This method helps prevent brief increases in amplitude to be classified as events. Moreover, it has shown to prevent underestimation of the duration of such transient single-trial events (Little et al., 2019). From the theta events we first estimated the theta event probability. Theta event % was estimated by binarizing the time periods in a trial where the theta amplitude was above the event length threshold and then averaging across trials within each condition.

Within a trial, three theta timings were estimated: the theta onset which is the time at which theta exceeds the event length threshold, the theta peak which is time corresponding to the peak of the theta event, and theta offset which is the time when the theta event falls below the threshold (also see Fig. 2b). For correct trials, theta events (i.e. the peak) that occurred before the response times were considered. If a trial had more than one theta event with peak times t1 and t2, it was split into two trials with events at time t_1_ and t_2_ respectively. These timings were then related to the timings of the conflict metrics, i.e., the partial error onset and the correct response onset within a participant using Pearson’s correlation (r) and partial correlation analysis.

**Figure 1.**
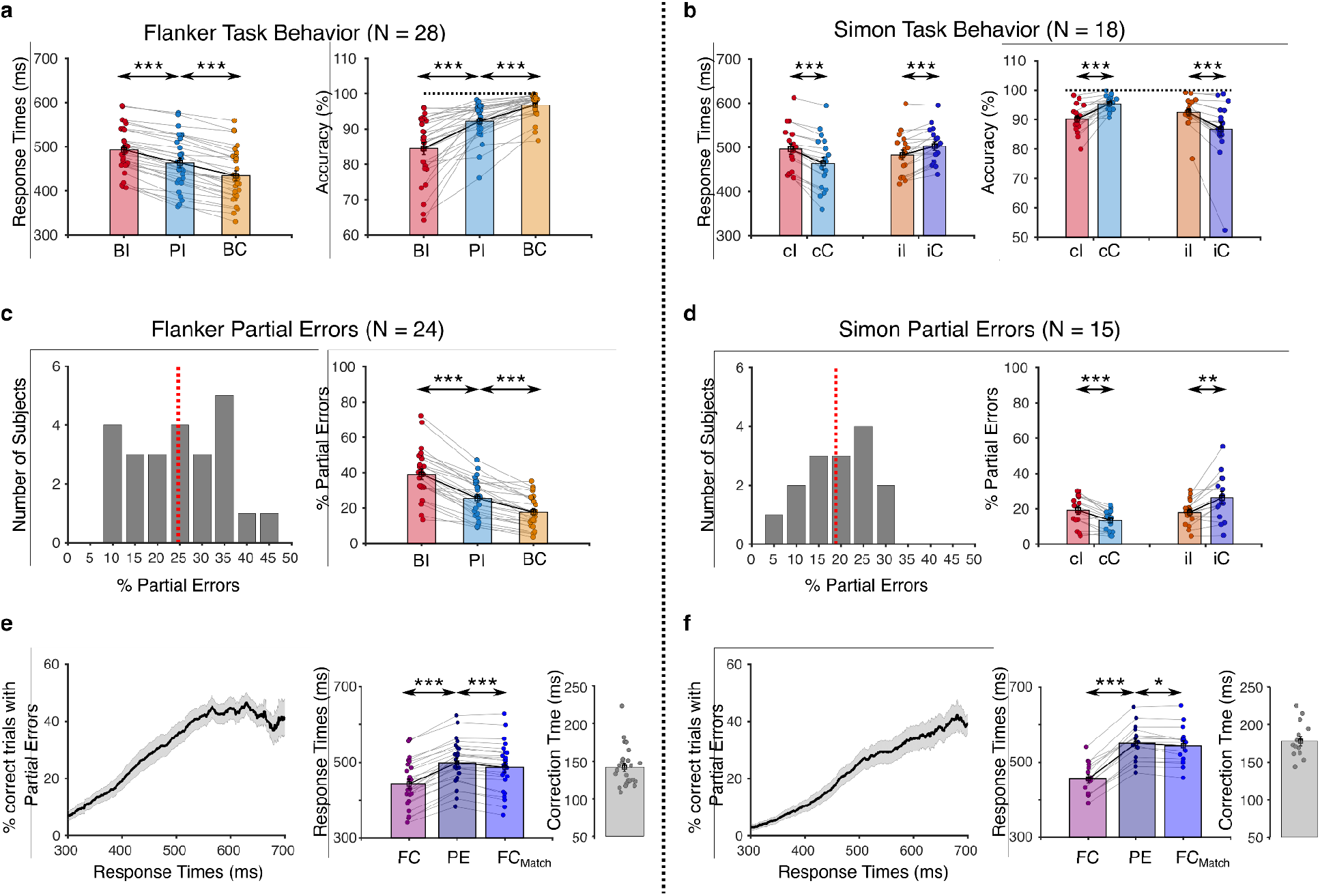
Behavioral data from the Flanker and Simon task indicate different levels of response conflict in different trial types. **a)** Response times and accuracies in the Flanker task are shown for the bilaterally incongruent (BI), partially incongruent (PI) and bilaterally congruent (BC) trial types. **b)** Response times and accuracies in the Simon task for the congruent and incongruent trials. Lower-case letters indicate whether the previous trial was congruent (c) or incongruent (i); upper case letters indicate the corresponding for the present trial. **c)** The left plot shows the percentage of trials with partial errors in each participant, with the red dotted line showing the average across all participants. The right plot shows the percentage of partial errors based on the trial type (dots indicate individual subjects, bars show average). **d)** Analysis corresponding to **(c)** is shown for the Simon task, with the right plot showing the proportion of partial errors as a function of previous trial conditions. **e)** Left plot shows for the Flanker task, the percentage of partial error trials as a function of response times. The middle plot shows the response times of all trials with no partial errors (i.e. full-correct trials, FC), of partial error trials (PE) and of trials matched for response time and stimulus type (full-correct matched trials, FCMatch). The left plot average correction time (dots indicate individual subjects, bars show average), which is time from the partial error EMG onset to the correct response EMG onset. **f)** The same analysis as **(e)** is shown for the Simon task.

**Figure 2.**
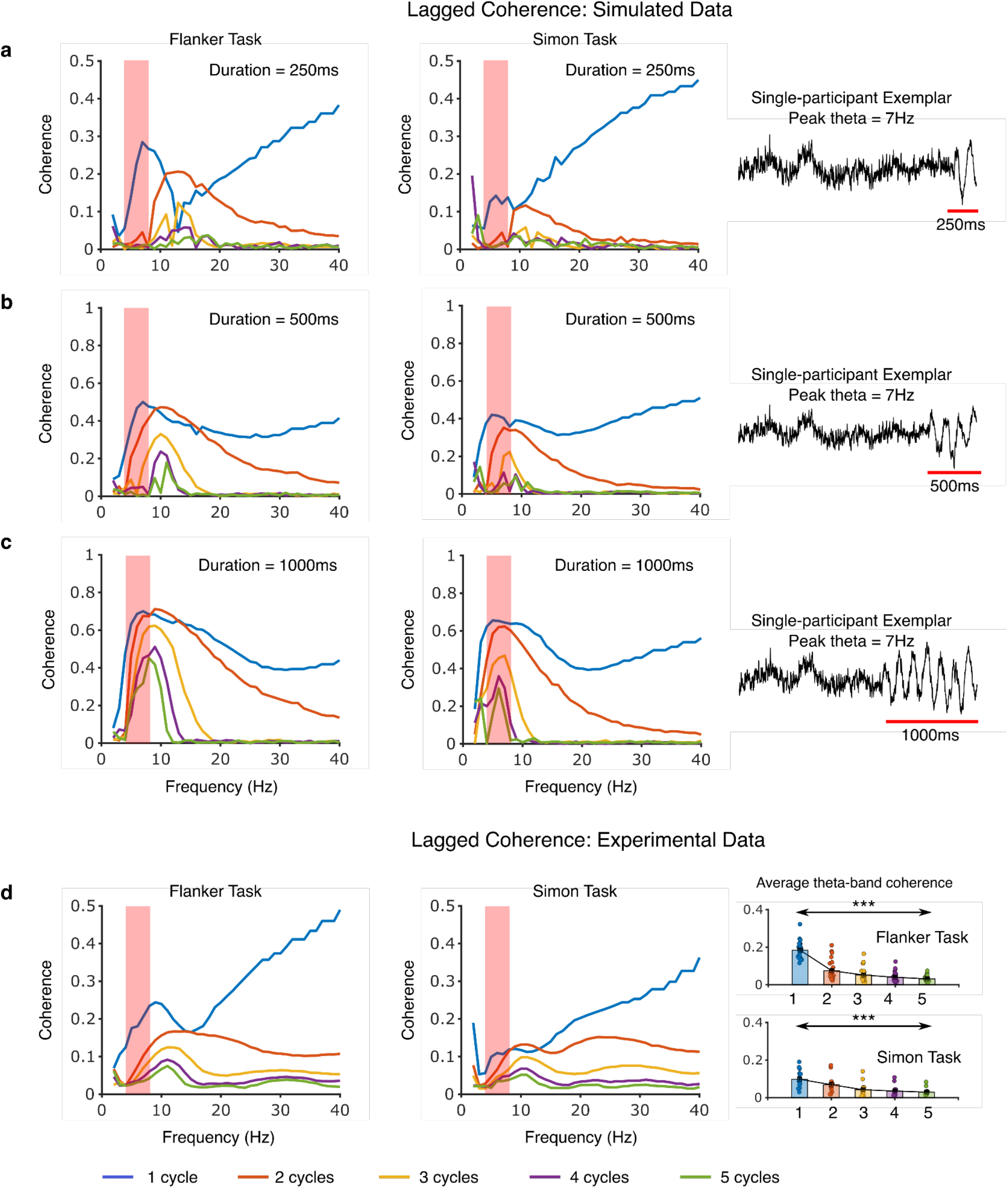
Lagged coherence reveals transient nature of mid-frontal theta. Averaged lagged coherence values for a simulated theta signal using the peak theta frequency for each participant for different durations **a**) 250ms, **b**) 500ms and **c**) 1000ms, computed for different window-lengths (i.e., cycle numbers: 1-5). The simulated signal contains an aperiodic part and an oscillatory part lasting for the duration as mentioned. The translucent red box denotes the theta-band frequency (4-8Hz). For a transient theta event, coherence is high and around the simulated frequency for 1 cycle. For longer theta signals the coherence becomes more concentrated around the simulated frequency for higher cycles. **d**) Pattern of average lagged coherence seen in the Experimental data (Flanker and Simon). The coherence at each cycle represents the average across all participants in that task (N = 24 in Flanker and N = 15 in Simon). The pattern in both the experiments show considerable similarity to the simulated scenario where theta was transient (i.e. the 250ms case). The bar-plots show that the average theta-band coherence across the different cycle numbers reduces a pattern also confirmatory of the fact that theta is transient in nature. Each dot represents a participant. The significance (***) represents results of a one-way repeated measures anova (see text for more details).

#### Lagged Coherence Analysis

In order to quantify whether the theta events detected from our amplitude thresholding methods in the both the tasks were indeed short-lived/transient events and not ongoing rhythms which were amplitude modulated, we computed lagged coherence. Lagged coherence is a metric that quantifies the rhythmicity of oscillations, unconfounded by amplitude modulations (Fransen, van Ede, & Maris, 2015). This forms an ideal metric to examine the temporal nature of the oscillations, i.e. transient or long-lasting. By computing the phase relationship between non-overlapping segments of a time-series signal, lagged coherence ascertains whether the phase is consistent across these time windows of interest. The size of the window is determined by the frequency of interest and the number of cycles of that frequency considered for computing the lagged coherence. For e.g. to compute the lagged coherence at 5Hz for 2 cycles, we would use a window length of 400ms. Thus, for a long-lasting rhythm, the phase will be better predicted across these time windows leading to larger coherence values compared to when the oscillations are transient. So, typically for bursts of oscillations, the lagged coherence overall would be smaller. Secondly, the lagged coherence would be higher around their frequency for shorter window sizes (expressed as the number of cycles of that oscillation, where phase can be better predicted across time-windows) and would decrease with increase in this window size.

In our study, we looked at the lagged coherence across all mid-frontal projected EEG data in each respective task. The lagged coherence analysis was performed using the NeuroDSP software for neural digital signal processing (Cole, Donoghue, Gao, & Voytek, 2019). For each participant, all trial epochs (partial error, full-correct and full-error) were concatenated to ensure there was a long enough time-window to estimate lagged coherence across all ranges of window sizes used to compute it. The coherence values were computed across a range of frequencies (2-40Hz) and window sizes (i.e. number of oscillatory cycles, 1-5). In order to understand the pattern seen in the experiment, we also simulated data having different durations of theta oscillations to see which pattern corresponds to the experimental data the best. For the simulated data, we initially created a 2.5-second epoch (comparable to the experimental epoch lengths) for each participant with every epoch containing a theta oscillation (at the participant’s corresponding peak theta frequency) of a specific duration (250ms, 500ms and 1000ms) and the rest was an aperiodic broadband noise which mimics the 1/f component of the background EEG. Furthermore, the signal to noise ratio was set at a factor of 4, i.e. the power of the theta was 4 times the level of the background noise at the same frequency. Similar to the experimental case, we simulated 1500 epochs (equivalent to the total number of trials in the experiments) which were then concatenated to then run the lagged coherence analysis. We ran the same analysis on these simulated data to see how the rhythmicity of theta affects the pattern of lagged coherence and compared it to the experimental data.

#### Gaussian Mixture Modeling of Theta Events

The theta timings (specifically the onset) were analyzed in relation to the partial error onset times using gaussian mixture modeling. We used this approach to dissociate the contribution of different theta modes, given that we observed bimodality in the shape of the difference distributions between theta onset and partial error times. The expectation maximization algorithm *(fitgmdist* function in MATLAB) was used to fit models with K gaussians. In our case K was set to either 1, 2 or 3. The covariance type was selected as full, with each component assuming any possible scale. We also set a 1% regularization to the data, with a maximum iteration of 1000. As a control condition, we also fit a generalized extreme value (GEV) distribution (*gevfit* function in MATLAB), which can model the location (mean), scale (variance) and shape (skewness) of the distribution as a potential model that could describe the underlying variability in theta timings. This was done to account for the fact that the underlying distributions of theta times could be still unimodal, but skewed. From the model fits, we obtained the Akaike Information Criterion (AIC) value and estimated the probability of each model (K = 1, 2 and 3 and GEV) as following 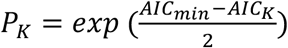. We then tested which of the models best predicted the underlying distribution of the theta timings. We also evaluated the mean and variance of the fitted gaussian models.

#### Fixed Effects Analysis: Joint Probability Distributions

The temporal structure of the theta activity was ascertained by computing the joint probability distribution of the theta onset times and the correct response EMG onset times. We pooled trials from both tasks to perform fixed-effects analysis. The probability value for a particular correct response and theta onset, say for instance at 300ms and 100ms respectively, was computed by first taking all the correct response onsets within 300 ± 25ms, and then looking at the number of trials that had a theta onset within 100 ± 25ms. This was then done for a range of correct response onsets (200 to 800ms) and theta event onset (−100 to 800ms). These probability maps were then thresholded for statistical significance by performing a permutation test against a null distribution which considered that the theta onsets were uniformly distributed across time (i.e. −100ms prior to stimulus to response time + 400ms) and corrected for multiple comparisons using false discovery rates (Benjamini & Hochberg, 1995). The same map for the trials in which a full error had occurred was computed differently. Since in these trials there are no correct response times, a pseudo “correct” RT was estimated by adding the individual’s mean correction time to the error response, i.e. the time from the partial error to the correct response onset. For the same trials, we considered those theta events which occurred between the stimulus and error response + participant-specific max correction time (i.e., average of the correction times greater than the 75th percentile). This was done to make sure we consider the events which occur even after the full error EMG onset, under the assumption that there could be temporal changes between trials where the error was partial compared to when it became a full error. This also ensures that the time window is long enough to detect any theta that might have occurred later, but within the potential time that would have been available to correct the error.

#### Statistical analyses

Data were analyzed using several different approaches, as appropriate for the specific comparison (detailed in the Results section) using the JASP software (JASP Team (2022). JASP (Version 0.16.3)[Computer software]). Some comparisons were performed using t-tests, either one sample or paired sample tests. The effect sizes for these tests were estimated using the Bayes factor in favor of the alternate hypothesis (small: BF_10_ = 1-3; medium: BF_10_ = 3-10; large: BF_10_ > 10). For distributions violating normality, we performed non-parametric Wilcoxon tests. For data with multiple levels, we used repeated measures (rm) ANOVA followed by post hoc t-test comparisons between specific conditions, corrected for multiple comparisons using Bonferroni correction. We reported the effect sizes for the ANOVA as partial eta square (η_p_^2^), interpreted as a small (0.01-0.06), medium (0.06-0.14) and large effect (>0.14). Data sphericity was controlled using the Greenhouse-Geisser correction. For the time-frequency analyses, the ERSPs, we used a non-parametric bootstrap method to compute regions of significance in the time-frequency space. We then used the false discovery rate to correct for multiple comparisons in these cases (Benjamini & Hochberg, 1995). Finally for the group-level comparisons of correlations we first used the Fisher-transform to convert the correlations to z values and then used rmANOVA and/or t-tests to compare between specific conditions. All data are presented as mean ± s.e.m.

## RESULTS

### Behavior

We analyzed behavior of subjects in the Flanker and Simon task by measuring RTs and accuracies in different trial types depending on the stimulus congruence. In a trial of the Flanker task the distractor stimuli on both sides could be different from the target stimulus (i.e. high conflict, bilaterally incongruent), the distractor stimuli on only one side could be different from the target stimulus (i.e. medium conflict, partially incongruent), or the distractor stimuli on both sides could be identical to the target stimulus (i.e. low conflict, bilaterally congruent). In line with previous observations, behavior from the Flanker task exhibited the expected patterns of slower RTs and decreased accuracy for incongruent compared to congruent trials (Appelbaum et al., 2011). The slowest RTs were observed in the bilaterally incongruent trials, followed by the partially incongruent and bilaterally congruent trials (F_1.5,39.2_ = 219.5, p < 0.001, η_p_^2^ = 0.9, Fig. 1a). The accuracy in each condition also scaled with the stimulus condition, with the lowest accuracy in the bilaterally incongruent trials (F_1.2,31.3_ = 50.9, p < 0.001, η_p_^2^ = 0.7).

In the Simon task, the stimulus could appear on the side of the fixation cross that is opposite to the correct response (high conflict, incongruent trial) or on the same side as the correct response (low conflict, congruent trial). Furthermore, behavior depends on the previous trial condition, i.e., the well-known sequence or the Gratton effect (Egner, 2007; Gratton, Coles, Sirevaag, Eriksen, & Donchin, 1988). Accordingly, conflict is increased when the congruence switches between two successive trials, which lead to slower RTs for both congruent-incongruent and incongruent-congruent trial sequences (Fig. 1b). In contrast, congruent-congruent and incongruent-incongruent trial sequences had faster RTs, reflected by a significant interaction (F_1,17_ = 48.5, p < 0.001, η_p_^2^ = 0.7). The same interaction effect was observed for the accuracies as well, with lower accuracy in the incongruent trials compared to the congruent, especially when the previous trial was congruent and vice versa (F_1,17_ = 26.9, p < 0.001, η_p_^2^ = 0.6). Therefore, behavior in both tasks was affected by conflict as expected (Appelbaum et al., 2011; Cohen & Ridderinkhof, 2013).

EEG analyses focused on partial error trials, defined as correct trials in which the incorrect (prepotent) response was partially activated (observed as a small burst in the EMG of the incorrect response hand), but did not develop into a full response and was followed by the correct response. In both the datasets, we identified partial error trials (see *Methods* for more details) and confirmed that their occurrence was as expected (Cohen & van Gaal, 2014). In four participants from the Flanker task and three participants from the Simon task noisy muscle activity led to sub-optimal detection of trials with partial errors and their onset timings and were therefore excluded, leaving 24 and 15 participants in each task, respectively. On average there were ~20% of trials with partial errors in both tasks (Fig. 1c and 1d). The percentage of partial errors scaled with the level of conflict, with more partial errors seen in the bilaterally incongruent trials in the Flanker task (F_1.1,26.1_ = 94.1, p < 0.001, η_p_^2^ = 0.8). In the Simon task, there was again a modulation based on previous trial condition, with more partial errors seen in incongruent-congruent and congruent-incongruent trial sequences compared to congruent-congruent and incongruent-incongruent sequences, leading to a significant interaction (F_1,14_ = 18.4, p < 0.001, η_p_^2^ = 0.6). The percentage of partial errors also increased with slower RTs (Fig. 1e and 1f). As a result, all full-correct trials, i.e. trials that were correct and did not have a partial error based on our algorithm, were faster on average compared to the partial error trials (Fig. 1e and 1f). To control for the fact that there were more full-correct trials than partial error trials and that they had faster RTs, we selected a subset of trials from the full-correct trials to match for both RTs and stimulus type, referred to as full-correct matched trials (see *Methods* and Fig. 1e and f). These trials were then used for comparison with the partial error trials in the EEG analyses below. Finally, we also estimated the correction time, which is the time taken from the onset of the partial error EMG to the onset of the correct response EMG. These typically ranged from 100-200ms (mean ± SEM: Flanker: 142 ± 6ms; Simon: 178 ± 6ms). Next, we sought to extract mid-frontal theta activity to understand the trial-by-trial dynamics and relationship to the mechanisms resolving response conflict.

### Midfrontal theta occurs as transient events within single trials

To identify and characterize single-trial mid-frontal theta events, we crafted a spatiotemporal filter that maximized midfrontal theta-band for each participant (Fig. 3a and see *Methods*). The EEG data were projected onto the selected filter and then temporally filtered at that individual’s peak theta frequency.

**Figure 3.**
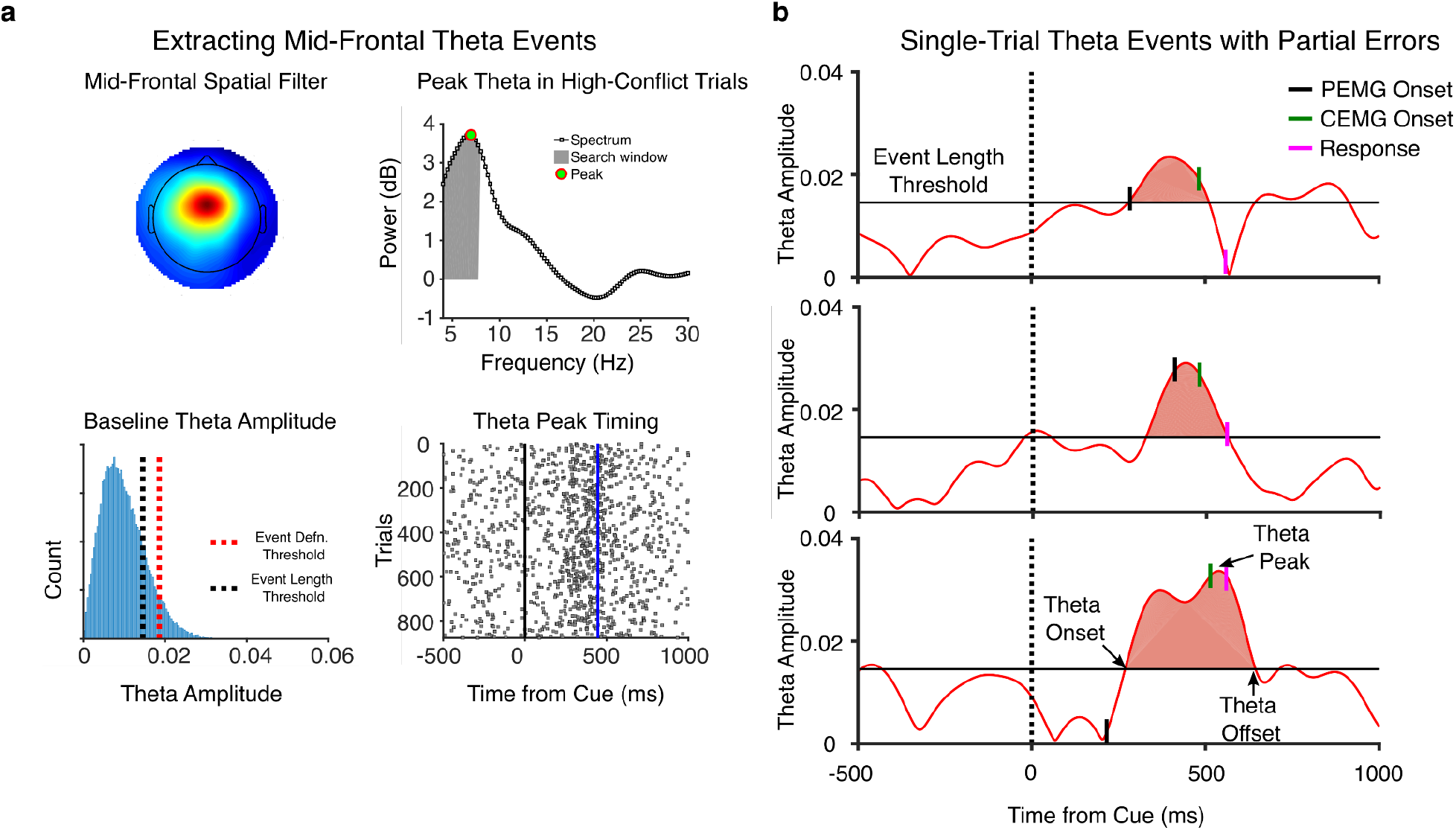
Theta Event Extraction. **a)** Shows the analysis steps for extracting theta events within a participant. Top left shows a mid-frontal spatial filter which was identified in an exemplar participant using GED in the Flanker task. The data were then projected on to the selected mid-frontal spatial filter. Top right shows the participant-specific peak theta estimated by looking at the normalized power spectrum of those trials capable of inducing response conflict the most, i.e., the bilaterally incongruent trials in the Flanker task and the incongruent trials in the Simon task (top right). All trials were filtered at this peak theta frequency using a frequency domain gaussian. Bottom left shows theta amplitude distribution from a baseline period (−500 to −100ms in relation to the stimulus) used to evaluate the event thresholds. To identify a theta event, median + 1.75SD was used as the threshold (red dotted line), and once an event was detected, its onset and offsets were determined by a lower threshold median + 1.0SD (black dotted line). Bottom right shows the raster plot of the timings of the theta peak for all trials, with the dotted blue line representing the mean response time across all trials. **b)** Example trials within a participant showing the theta event, its timing (onset, peak and offset) in relation to the timing of the conflict related muscle activity, the partial error onset (PEMG), the correct response onset (CEMG) and the response time.

Before extracting single-trial theta events using our amplitude threshold approach (see *Methods*), we characterized the dynamics of theta in both tasks. The goal was to determine whether theta oscillations were mostly ongoing and modulated in amplitude by task events, or whether they predominantly occurred as short-lived bursts of activity. To do so we used a measure called lagged coherence (Fransen et al., 2015), a metric reflecting the rhythmicity of underlying oscillatory activity. Lagged coherence quantifies phase predictability across non-overlapping time windows, so that a transient oscillation will have lower lagged coherence value compared to an ongoing rhythm. In the analysis window sizes are systematically varied in length (in units of oscillations cycles), and for longer time windows the coherence will decrease strongly if the length of the oscillations is shorter than the time window.

To confirm that the lagged coherence measure would allow us to distinguish between longer, ongoing theta and brief theta bursts, we first generated simulated data for all participants using their peak theta frequency (see exemplar simulated time series in Fig. 2a, b and c). For each participant we simulated a theta oscillation of varying duration, lasting 250ms, 500ms or 1000ms (Fig. 2a, b and c, see Methods for details) and then calculated the lagged coherence for a range of frequencies (2-40Hz) and numbers of cycles (1-5).

Simulating a transient theta signal (250ms, Fig. 2a) for the participants of both tasks lead to a pattern of lagged coherence with increased coherence in the theta band only for a very short time window (length of 1 cycle), which then decreased and shifted in frequency as the window size (i.e. number of cycles, colored lines in Fig. 2) increased. This pattern was different when the theta was longer lasting (i.e. 500ms and 1000ms, Fig. 2b and c) with a coherence peak in the theta band also for longer time windows. Comparing this to the experimental data from the Flanker and Simon task, in which we computed lagged coherence across all participants, we observed a pattern similar to the simulations with a short-lived theta (i.e. 250ms, Fig. 2d). Again, the lagged coherence peaked for a window size of 1 cycle, and for longer time windows the coherence values decreased in the theta band and the coherence peak shifted towards higher frequencies (Fig. 2d). A one-way repeated measures ANOVA revealed a main effect of window length on the lagged coherence in the theta-band (4-8Hz) as it decreased with increasing window length for both tasks (Flanker: F_2,47_ = 114.2, p < 0.001, η_p_^2^ = 0.9; Simon: F_2,27.7_ = 43.0, p < 0.001, η_p_^2^ = 0.8), supporting a transient nature of the underlying theta. We next used our amplitude-threshold method to extract these transient single-trial theta events.

Using the median theta amplitude in a baseline period prior to the stimulus, a threshold (event definition threshold, median + 1.75SD) on the theta baseline amplitude was defined. Using this threshold, peaks of theta activity that exceeded this threshold were identified. From the identified peaks, using a lower threshold (or event length threshold, median + 1SD), a theta “event” was defined as any time period where the theta amplitude was above this threshold (Fig. 3a, see *Methods* for more details). Within a participant, there was both an increase in the number of theta events and variability in the timing of these events in and around the time of response (Fig. 3a, bottom left). Furthermore, the duration of these theta events irrespective of the stimulus condition or time window were on average around 200-300ms (Flanker: 229 ± 6ms, Simon: 259 ± 4ms) pointing towards their transient nature and corroborating our findings from the lagged coherence analysis on the transient nature of these events. We also observed a high correlation between mean theta amplitude and the number of theta events in a trial at our event definition threshold (see Supplementary Information Fig. S1), an approach taken to justify threshold selection as seen in previous studies looking at beta bursts (Little et al., 2019).

### Theta event probability increases in Partial Error trials

The classic event-related spectral perturbation analysis (ERSPs) for all conditions of the mid-frontal projected data revealed the typical increase in theta power for both tasks (Fig. 4a). Furthermore, a comparison of the theta power increase in the full-correct matched trials and partial error trials confirmed higher theta power in partial error trials in both tasks, which is an observation from earlier studies (Cohen & van Gaal, 2014). As shown before, at a single-trial level we found that theta oscillations manifested as transient events and could be characterized by an onset, a peak and an offset (partial error trials shown in Fig. 3b).

**Figure 4.**
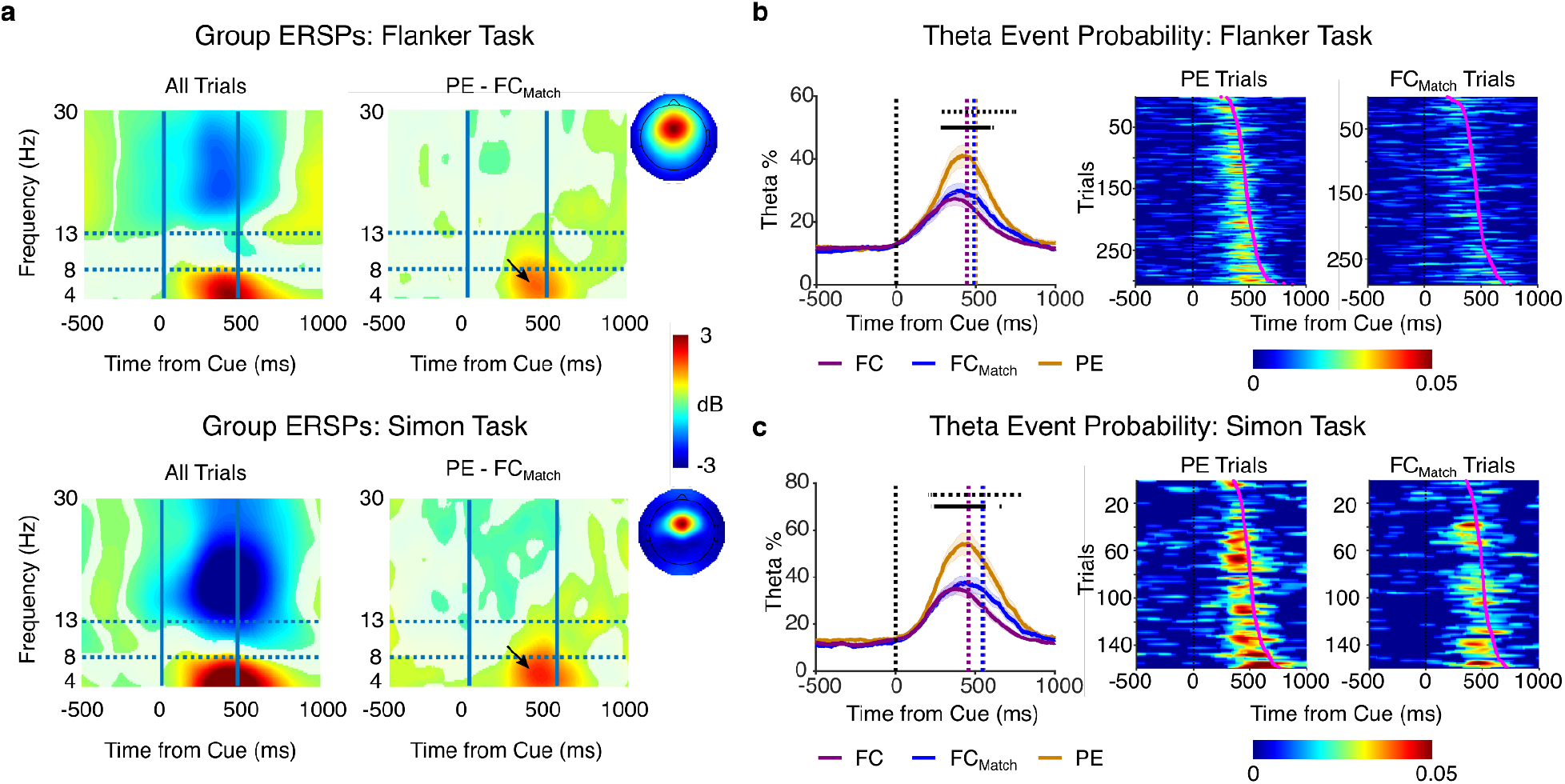
Theta Event % in Partial Error and Full-correct trials. **a)** Group-level ERSPs (p <0.05 FDR corrected maps) for both tasks, showing all trials and the difference between the partial error and full-correct matched trials. The arrow represents the region of theta power difference seen in both tasks for the PE-FC_Match_ ERSPs. **b)** The left plot shows theta event % seen in the partial error trials compared to all full-correct and the matched trials in the Flanker task. The horizontal lines show the regions of significant difference using a permutation-based t-test between the partial error and full-correct trials (dotted black line) and partial error and full-correct matched trials (solid black line). The middle and right plots are single subject examples of theta events sorted by response time. **c)** Same as (**b**) but for the Simon task.

The increase in theta power seen in the partial error trials (Fig. 4a) suggests that a potential reason for this observation could be the increase in the number of theta events in the partial error trials compared to the full-correct trials. We examined how the number of theta events is modulated over time and found a significant increase in the frequency of theta events in the partial error trials compared to both full-correct and full-correct matched trials in both task paradigms (Fig. 4b). This is confirmed by the fact that there were more trials containing at least one theta event in the partial error trials compared to the full-correct matched trials (Flanker: 54 ± 3% vs 44 ± 3%; t_1,23_ = 6.1, p < 0.001, BF_10_ > 100 and Simon: 67 ± 4% vs 53 ± 3%; t_1,14_ = 8.4, p < 0.001, BF_10_ > 100). This was also visible in single participants suggesting that some theta activity could have been evoked by the partial errors. However, the change in theta power could also be driven by other factors such as higher amplitude events in the case of the partial errors or a different underlying temporal distribution of theta events in the partial errors compared to the full-correct matched trials. This highlights the utility of decomposing theta into single-trial events in providing a better characterization of neural activity occurring during both trial scenarios in comparison to just oscillatory power. Next, we investigated how the timing of these single-trial theta events relate to the conflict-related EMG timings.

### Theta event onset times reveal two modes of mid-frontal theta activity

For trials in which theta events occurred between the stimulus onset and the response time, we estimated the time (onset, peak and offset) when these events occurred in relation to the EMG signals of partial errors and correct responses. Given that we had already established that theta is short-lived, we decided to look at theta onset times as they would be informative about when conflict-related processing began. In the partial error trials, within participants there was considerable variability in the timing of theta activity and we noticed that the distribution of theta onset times relative to the partial error times was typically bimodal (Fig. 5a and b). Using gaussian mixture modeling with K = 1, 2 and 3 number of gaussians, we probed which model best captured individual distributions (see *Methods* for details). We found that the model with two gaussian components fitted the distributions better than one (Fig. 5a and b top left; Flanker: W = 32.0, p < 0.001, BF_10_ > 100; Simon: W = 16.0, p = 0.010, BF_10_ = 16.0). However, this increase in the model probability was not just due to increase in the parameters, as K = 3 did not improve the fit in comparison to K = 2 (Flanker: W = 173.0, p = 0.527, BF_10_ = 0.2; Simon: W = 79.0, p = 0.303, BF_10_ = 0.4). As an additional control, we also fit the data to a generalized extreme value (GEV) distribution, as it can model location, scale and shape of the underlying distribution. This was done for two reasons: firstly, a K=1 gaussian model could naturally be worse in fitting the data if there is skewness in the distributions (which we observed for some participants; see Supplementary Information Fig. S2–5 for individual distributions and the fits for the GMM and the GEV models) which the GEV can capture, and secondly the GEV is well-suited to capture distributions which have a sharp cutoff which is a possibility given that we look at theta events within a fixed window of time (i.e. stimulus to response time). We still found that the K=2 model was the more probable model in comparison to the GEV (Fig. 5a and 5b; Flanker: W = 249.0, p = 0.004, BF_10_ = 42.0; Simon: W = 105.0, p = 0.008, BF_10_ = 14.7). All these findings point to the idea that there are two modes of theta activity in partial error trials.

**Figure 5.**
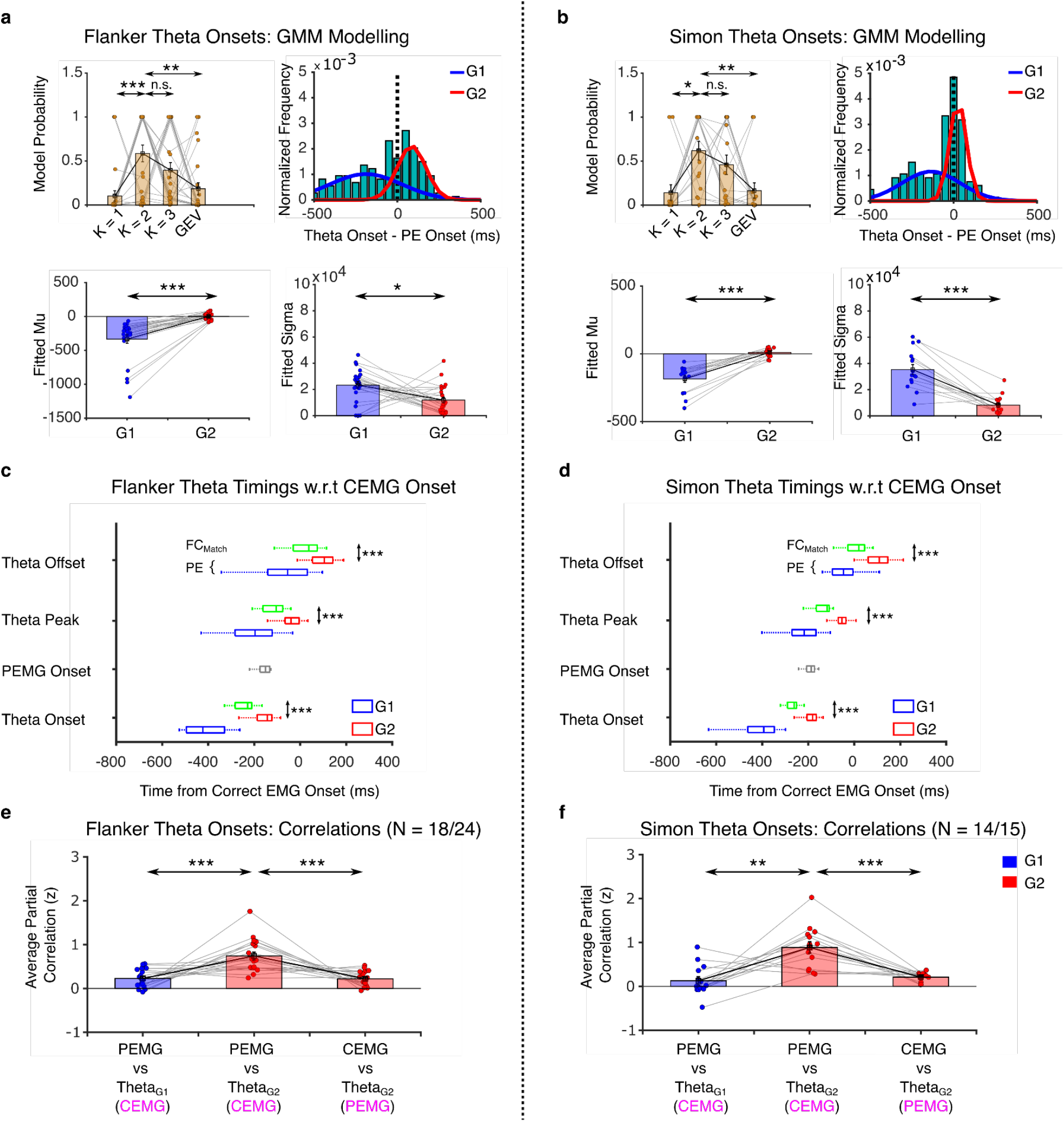
Different modes of theta activity. **a)** Top left plot shows the model probability for the gaussian mixture modeling (GMM) with K = 1, 2 and 3 gaussian components and the generalized extreme value distribution (GEV) in the Flanker task in the partial error trials. Top right shows a single subject exemplar for the distribution of theta event onsets in relation to the partial error onsets and the gaussians fitted using K = 2 model. Bottom left and right plots show the fitted parameters mu and sigma for both the gaussians respectively. The parameters suggest that there are two modes of theta, with one peaking prior to the partial error (G1) and the other peaking around it (G2). **b)** Same as **(a)** but for the Simon task. **c)** and **d)** Boxplots showing the mean theta event timings in the partial error (PE) and full-correct matched trials (FC_Match_, green box). For the partial error trials, the theta timings have been split into those belonging to G1 (blue box) and G2 (red box). **e)** and **f)** Fisher-transformed partial correlations between the theta onsets belonging to either G1 or G2 and the EMG timings (partial error – PEMG and correct response – CEMG) in the partial error trials for the Flanker and Simon task respectively. The magenta colored variable is the control variable used in the computation of the partial correlation in each case. Analysis shows that the relationship is strongest for the theta onset times belonging to G2 and the partial error onset, suggesting that one mode of theta is strongly linked to the partial error.

Next, to see whether this bimodality in the theta onset times was specific to the partial error trials, we investigated the full-correct matched trials as well. We assigned a surrogate “partial error onset” time to each full-correct trial, which was obtained from the partial error trial that was the closest match to the full-correct trial in terms of the response time. Performing the gaussian mixture modeling on this set of trials, the evidence that a bimodal distribution of theta onset times exists in these trials was not strong as in the case of the partial error trials. In the Flanker task, although there was a significant increase in the model probability for K = 2 compared to K = 1, the Bayes factor evidence was medium (BF_10_ = 4.5; 0.27 ± 0.08 vs 0.55 ± 0.09; W = 73.0, p = 0.027). In the case of the Simon task there was no significant difference between K = 1 and K = 2 with weak Bayes factor evidence in support a bimodality (0.25 ± 0.10 vs 0.48 ± 0.11, W = 33.0, p = 0.135, BF_10_ = 0.9). Furthermore, there was no evidence that the K=2 mixture model fits better than the unimodal GEV model in both tasks (Flanker: 0.55 ± 0.09 vs 0.44 ± 0.09; W = 188.0, p = 0.290, BF_10_ = 0.5 and Simon: 0.48 ± 0.11 vs 0.62 ± 0.11; W = 39.0, p = 0.252, BF_10_ = 0.4). The same result was observed when, instead of the surrogate partial errors, we looked at the theta onset times aligned around the correct EMG onsets, i.e. an actual task event. This presents convincing evidence that a bimodality does not exist in the full-correct matched trials, suggesting that the two modes seen in the partial error trials might have different functional roles.

To probe this, for each participant, using the fitted parameters from the K = 2 model, we obtained a mean μ and a variance *σ* for each gaussian in the partial error trials. The first mode occurred prior to the partial error onset (**G1** mean ± SEM, Flanker: −333 ± 62ms and Simon: −185 ±26ms), while the second mode occurred around the partial error onset (**G2** mean ± SEM, Flanker: 2 ± 10ms and Simon: 11 ± 9ms). In addition, G2 had a smaller variability than G1 in both tasks (Fig. 5a and 5b bottom right), in line with interpretations tying the G2 theta events to the possible error that could have been committed. As the exact timing of the G2 theta events relative to the partial errors possibly constrains potential functions related to error monitoring or correcting, we next examined the different theta modes in more detail.

We grouped theta events into G1 and G2 events by comparing the probability density functions of the two fitted gaussians at a given theta onset time. The theta event was then assigned to either G1 or G2 based on which had the higher relative likelihood. We initially looked at the timing of the theta events in the partial error and full-correct matched trials in relation to the correct response EMG onset (Fig. 5c and 5d), but plotted the theta times in the partial error trials split into G1 and G2. As expected, the G2 theta onsets were present closely around the time of the partial error onsets, pointing to their role in processing the error, with the G1 theta onsets happening much earlier, suggesting their role in potentially early stimulus conflict processing. Interestingly, the theta onsets (and as a result the peak and offset times) in the full-correct matched trials happened earlier compared to the G2 theta (Both tasks: t > 6, p < 0.001 and BF_10_ > 100) which links the G2 theta strongly to the partial errors, potentially involved in correcting the impending error.

In order to establish this connection, we correlated the theta onset times belonging to G1 and G2 to the partial error EMG timings within each participant. Here, we used partial correlations as any relationship between the theta timings and the EMG timings could be mediated by the correlations between the partial error EMG onsets and the correct response EMG onsets, and we would have to control for this effect. So, when correlating theta timings from G1 and G2 to the partial error EMG timings, the correct response EMG timings were used as the control variable. This was vice versa when the theta timings were correlated to the correct EMG onsets. We found that the partial error onsets correlated stronger with the G2 theta onset times compared to G1 (Fig. 5c and 5d; Flanker: t_1,17_ = 4.0, p < 0.001, BF_10_ = 42.1; Simon: t_1,13_ = 4.1, p = 0.001, BF_10_ = 31.5). Furthermore, the correlation of G2 with partial errors was stronger than the correlation of G2 with correct response onsets (Fig. 5c and 5d; Flanker: t_1,17_ = 4.3, p < 0.001, BF_10_ = 68.8; Simon: t_1,13_ = 4.9, p < 0.001, BF_10_ > 100). This supports that theta activity can be linked to partial errors. Due to the mixture model fitting, not all participants had a sufficient number of trials for this correlation analyses in either G1 or G2. Therefore, we present results of participants who had 15 or more data points (i.e., theta onsets) in both G1 and G2, which led to having 18/24 and 14/15 participants in the Flanker and Simon task, respectively. To see whether the results depended on the specific inclusion criteria, we repeated the analysis and included participants with 5 or more data points, leading to 20/24 and 15/15 participants from the Flanker and the Simon task, respectively. The relationship between theta timings and the EMG timings remained the same for both task scenarios (see Supplementary Information Fig. S6).

In addition, for these participants we examined the duration and amplitude of theta events in partial error trials. We found that the theta event durations from G2 were smaller compared to those from G1 (Flanker: 309 ± 13ms vs 255 ± 8ms; t1,17 = 5.0, p < 0.001, BF_10_ > 100; Simon: 352 ± 10ms vs 296 ± 11ms; t_1,13_ = 5.0, p < 0.001, BF_10_ > 100). The theta amplitude, however, was stronger for G2 theta events compared to G1 (Flanker: 0.032 ± 0.002 vs 0.034 ± 0.002; W = 19.0, p = 0.002, BF10 = 80.5; Simon: 0.039 ± 0.001 vs 0.042 ± 0.001; W = 9.0, p = 0.004, BF_10_ = 25.5), suggesting a strong and punctate activity of G2 theta in response to the error. These results support the idea that there are several modes of theta activity that are likely driven by different events. The first mode (captured by G1) is possibly related to the level of detected conflict, while the second mode (captured by G2) may be more directly linked to the processing of the partial error.

### Temporal structure of theta activity suggests possible role in error-correction

The relationship between one of the theta modes (i.e. from G2) and the onset of partial errors suggested that this theta could be a neural signature of overcoming the potential error, i.e. error-correction. To investigate this, we performed a fixed effects analysis where we separately pooled all the partial error trials from all subjects across both tasks (given that all of the theta event results were the same across both the tasks). We then looked at the joint probability distribution of the theta onset times in these trials as a function of the correct response EMG onset. The goal was to obtain the underlying temporal structure of theta across time that was yielding the correlations analyzed above (Fig. 5e and 5f). To examine a potential role of theta in the correction of errors, we also investigated trials in which the participants made a response error, i.e. the full error trials. In these trials, we analyzed theta events that happened in between the cue onset and the error response plus the participant-specific max correction time. The rationale was that if theta events reflect a process that prevents errors, they would have been evoked within this time period (see *Methods* for more details). Since there were only a small percentage of error trials per participant, we decided to combine the data from both tasks here as well. We again performed a gaussian mixture modeling analysis on theta onsets in the error trials. We kept the model parameters the same as for the partial error trials, i.e. K=2.

We found that the theta activity from G1 in the partial error trials was concentrated prior to the partial error onsets (cyan dots in Fig. 6a). In contrast, the theta events from G2 were distributed around the partial error time onsets, following them in orientation (Fig. 6b). This was evident across the whole temporal range of correct response onsets. The tight concentration of G2 theta events around the partial errors in partial error trials points to a potential role of theta in the correction of the error that has already started to activate the muscles.

**Figure 6.**
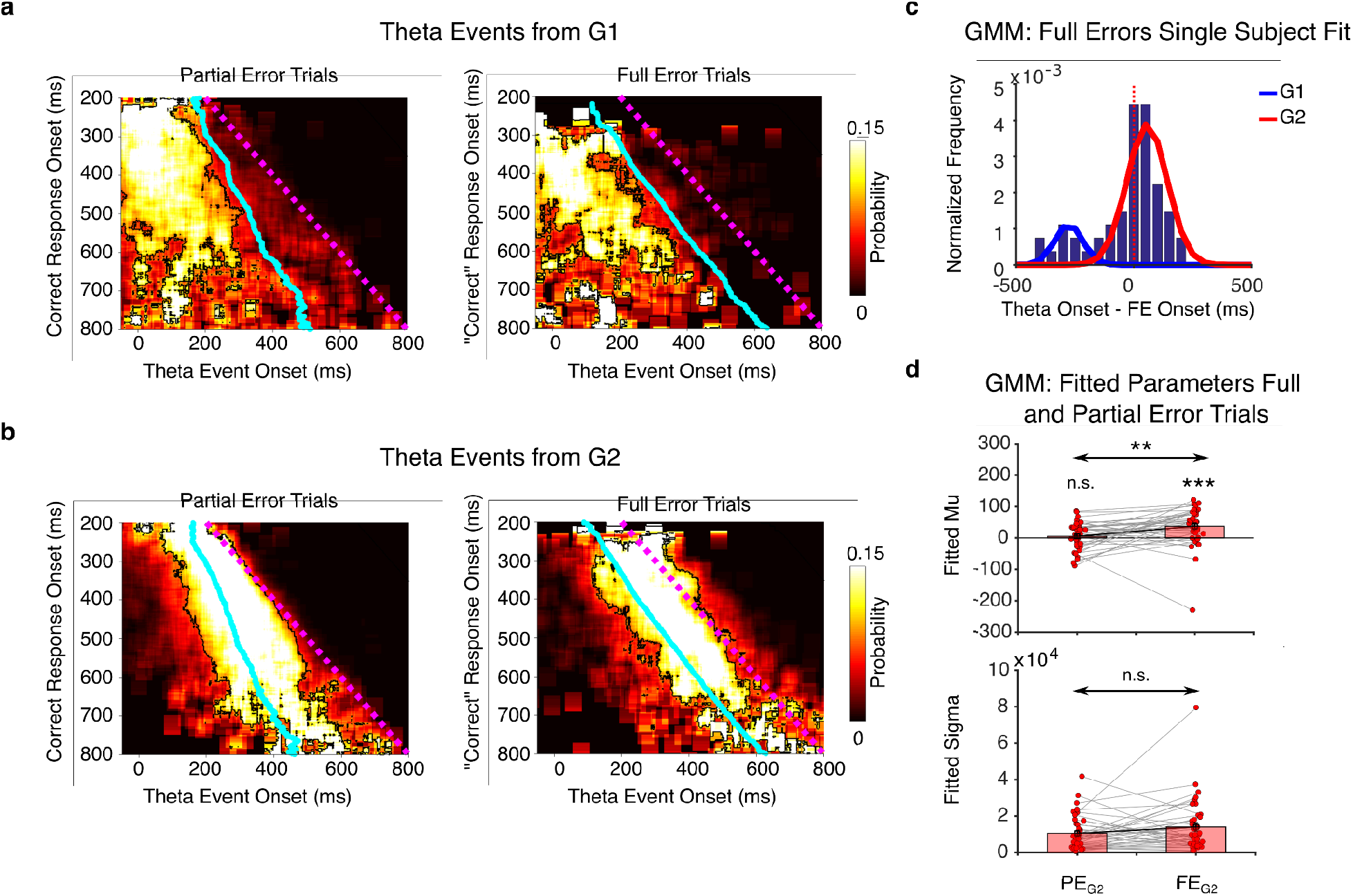
Temporal structure of theta onsets. **a)** The joint probability distribution of theta onsets from the G1 component of the GMM fit and the correct response onsets (dotted magenta line indicates where theta onset equals correct response onset) for the partial error and full error trials. **b**) Same as (**a**) but for the theta onsets belonging to the G2 component. In both (**a**) and (**b**), the cyan line indicates the average partial error onset times for the corresponding correct response onsets. For the full error trials, the cyan line indicates the full error EMG onset times and the magenta line corresponds to the potential “correct” response obtained by adding the participant-specific mean correction time.The contoured regions represent fdr corrected significance maps, showing increase in probability in comparison to a null distribution assuming that theta onsets were uniformly distributed. **c**) Single subject exemplar of the theta onset distributions for the full error trials with G1 and G2 components superimposed. **d**) The gaussian mixture model parameters (mu and sigma) for the full error and partial error trials for both tasks, specifically for the theta onsets from G2. Each dot represents participants from both tasks.

Similar to the partial error trials, full error trials also had a mode prior to the full error onset (G1) and another mode in and around the full error onset (G2, single subject example shown in Fig. 6c). In the temporal structure of the full error theta onsets from G2, the probability distribution aligned well with the full error onsets, but appeared to be shifted in time compared to how theta aligned in partial error trials (Fig. 6b). This shift was evident in the parameters of the gaussian fits with the average theta onsets from G2 being significantly later than the full error onsets (Fig. 6d, 37 ± 10ms, t_1,38_ = 3.7, p < 0.001, BF_10_ = 43.2), and more importantly their timing was significantly later than the G2 theta onsets in the partial error trials (37 ± 10ms vs 5 ± 7ms, t_1,38_ = 2.9, p = 0.007, BF_10_ = 5.6). Moreover, the variability of the G2 distributions from the full error and partial error trials did not change, suggesting that they could be the same process which just got engaged a bit too late in the full error trials (W = 293.0, p = 0.180, BF_10_ = 0.5). This supports the idea that the theta related to the partial error might be involved in error-correction.

## DISCUSSION

In this study we investigated the relationship between single-trial theta events and metrics of response conflict behavior. We analyzed neural data from the well-known Flanker and Simon tasks known to induce response conflict. Average increases in theta commonly seen during these tasks appeared as short transient increases in activity, akin to bursts of oscillations typically seen in other domains of motor control, for instance beta bursts and motor inhibition (Errington, Woodman, & Schall, 2020; Jana, Hannah, Muralidharan, & Aron, 2020; Muralidharan et al., 2021; Wessel, 2020). This transient activity, which we termed “theta events,” was variable in both its timing and occurrence based on the trial type with different degrees of conflict.

Specifically, by investigating those trials in which an error was partially committed, as identified from the muscle activity of the incorrect response, we showed that these theta events were increased compared to matched trials without partial errors, indicating that this theta could be evoked by potential, upcoming errors. The timing of these events in the partial error trials showed that there could be multiple theta components, with one of them occurring prior to the partial errors and the other in and around the time of a partial error. However, on closer inspection in single trials, our analyses revealed two modes of theta, one of which was strongly related to the partial error onsets. This implies that there are different forms of theta activity linked to the underlying processing of conflicts and potential errors.

### Theta timing points to a role in error-correction

Most studies on theta and conflict-resolution focus on the relationship between theta power and RTs. However, in our study decomposing theta activity into single-trial events revealed finer temporal relationships, specifically to conflict- and error-related processing. From the identified two modes of theta, one mode (theta events from G1) was weakly modulated by the error and could have been evoked by the conflict detected at the level of the stimulus given its presence prior to the partial errors. Once conflict was detected, the rapid engagement of theta could be sufficient in most cases to prevent errors in the full-correct trials. On the other hand, in the partial error trials, this mode of theta could still have been evoked as a result of stimulus processing but perhaps not enough to prevent the error from eliciting a muscle response. For clarity we refer to this theta here as “conflict-related theta”. The other mode (theta events from G2) seemed to strongly track the timing of partial errors, suggesting that this theta could also have been elicited in reaction to a potential, upcoming error. Possible functions of this mode of theta include the detection, the monitoring, and even the correction of the error, so we refer to it as “error-related theta”. This error-related theta would then reflect the cognitive processes aiming to prevent an error from releasing completely.

Evidence that theta activity represents neural processing involved in error-correction has been previously hinted at, with greater theta power correlating to shorter correction times in partial error trials (Cohen & van Gaal, 2014). From our study, two pieces of evidence add to the proposed error-correcting role of theta. Firstly, the stronger relationship between theta activity from this mode and partial error onsets revealed a different temporal structure (i.e. presence of a bimodality) of theta activity than in full-correct trials. Secondly, the timing of the error-related theta occurred a bit later in the full error trials compared to the partial error trials. Theta activity over different time-scales has been shown to interact with each other to help in online behavioral adaptation (Cohen, 2016). It could be that the error-related theta events we observed in the data, and their short-timescale, could be involved in the rapid correction of the partial error, and thus if they occur a bit later in time, this leads to a full error.

### Theta modes might have different underlying sources and relate to different cognitive strategies

Results of our study suggest that the modes of theta we observe seem to represent processes involved in conflict and error-processing. Previous studies have shown that both conflict and error-related processing could be functionally dissociated both in terms of the areas they originate from and the frequency bands in which they operate (Cohen & van Gaal, 2014; Nee, Kastner, & Brown, 2011; Ullsperger & von Cramon, 2001). In our case, these modes of theta also differ in terms of their strength and duration, with the error-related theta events being stronger and more punctate, compared to the conflict-related theta. The dissociation with respect to their functional role and properties could reflect that these two modes of theta are generated by different underlying brain networks. This is supported by studies that have demonstrated the presence of multiple sources of mid-frontal theta (Töllner et al., 2017; Zuure, Hinkley, Tiesinga, Nagarajan, & Cohen, 2020), and related them to both conflict and other conflict-independent processes (Mückschel, Dippel, & Beste, 2017; Töllner et al., 2017). In addition, our study provides a detailed picture of the temporal dynamics of theta and their potential role in correcting potential erroneous responses. The findings that there are multiple modes of theta which are temporally separable has implications as being targets for brain-stimulation studies to verify the functional dissociation between these two modes of theta.

If the two theta modes indeed reflect two different functions in the processing of conflicts and errors, it seems plausible that there is also some individual variation in how these two systems are engaged when dealing with conflict. For example, we noticed that for a minority of participants a single gaussian provided a better fit than two gaussians in the gaussian mixture modeling of the theta onsets in the partial error trials (Fig. 5a and 5b). Thus, it is possible that this was due to exerting a particular control strategy, such as initiating theta activity proactively to prevent errors from occuring, or relying more on the error-correcting mode of theta instead. Although mid-frontal theta is considered to be a general mechanism activated whenever there is a need for control (Cavanagh & Frank, 2014; Cavanagh & Shackman, 2015; Cohen & Donner, 2013), theta has been also implicated in different control strategies (Cooper et al., 2015; Eisma, Rawls, Long, Mach, & Lamm, 2021; Messel, Raud, Hoff, Stubberud, & Huster, 2021), such as proactive or reactive (Eisma et al., 2021). While the present data did not allow us to test these ideas, it points to potentially interesting future studies examining how different degrees and strategies of cognitive control affect theta signatures. For example, one prediction would be that a shift in the cognitive strategy is accompanied by a shift between the two modes of theta.

### Models of theta function and conclusions

Our findings have implications for rise-to-threshold models incorporating theta activity, specifically the interpretation that theta activity affects decision/response thresholds or “hold your horses” forms of control (Cavanagh & Frank, 2014; Cavanagh et al., 2011; Zavala et al., 2013). For example, if the function of both modes of theta involves increasing decision thresholds, important questions arise on how the error-correction would manifest mechanistically, and what the mechanistic differences are between the two modes. One possibility is that they differ in terms of their selectivity, so that e.g. the error-related theta raises thresholds selectively for the incorrect response. To the contrary, there is evidence that cortico-spinal excitability is decreased during conflict resolution (Duque, Olivier, & Rushworth, 2013; Wessel, Waller, & Greenlee, 2019), which is non-selective to the action effector, suggesting a global mechanism at play to resolve competition between responses. However, these studies have not looked at partial error trials or the possibility of theta having different modes. Therefore, it is possible that the conflict-related theta might reflect a “caution” signal, globally holding responses from execution, and the error-related theta represents processes more specific to the incorrect response. Future studies aiming to resolve this would further help in understanding theta’s role in error-processing better.

Alternatively, there could be other mechanisms implementing error-correction apart from rise-to-threshold model interpretations. For instance, theta’s interaction with downstream subcortical regions, specifically the basal ganglia, could be relevant for the different theta modes (Zavala et al., 2013, 2016, 2017, 2018). Our results point to the importance of the timing of the error-related theta, as its onset was overall delayed in the full error trials compared to the partial error trials. This could mean that how cortical theta, specifically the error-related theta, interacts with subcortical regions (e.g. the subthalamic nucleus) might drive successful error correction.

To conclude, we investigated the single-trial relationship between transient theta events and muscle-level metrics of response conflict. We showed that there is evidence for two modes of theta, one of which is strongly linked to the potential upcoming error. Future studies might further address the origin of the theta events and their interaction with neural circuits involved in error processing.

## Acknowledgements

This work was supported by the National Institutes of Health (DA026452 and NS106822). MXC was funded by an ERC-StG 638589 and a Hypatia fellowship from the Radboud University Medical Center. RS was supported by the EU Horizon 2020 programme through the FET Flagship Human Brain Project (HBP-SGA3, 945539).

## Supplementary Information

### Threshold selection for theta event detection

**Fig. S1.**
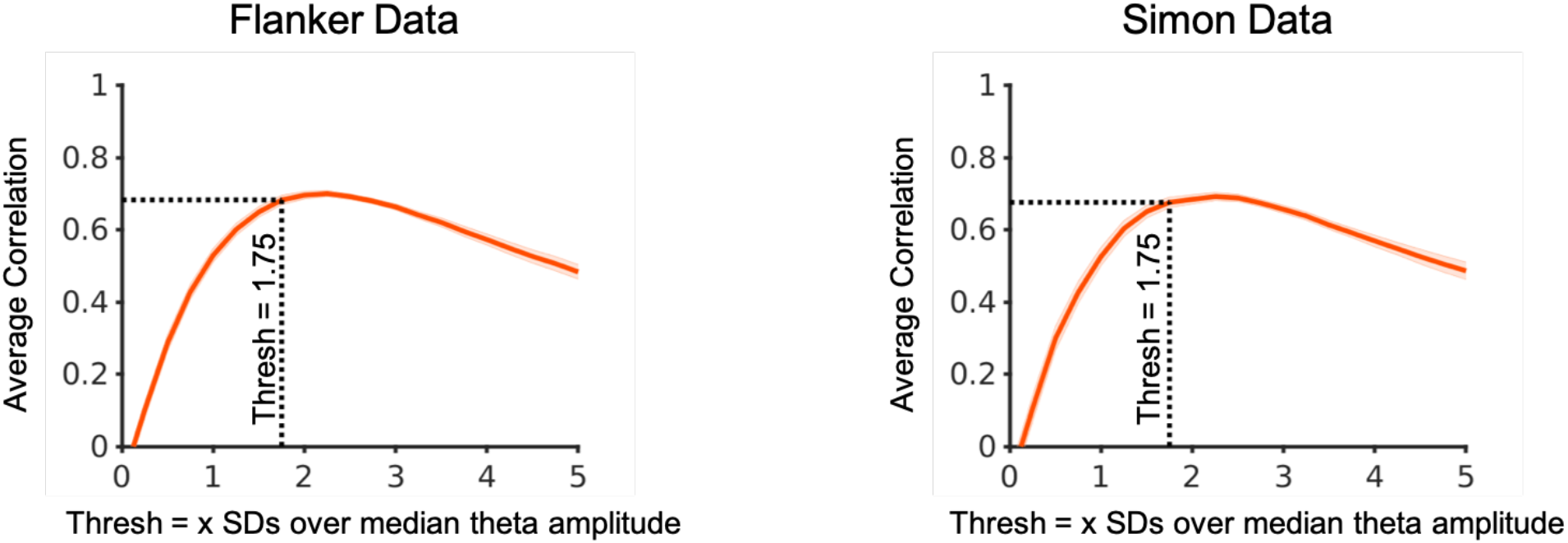
Average correlation between mean theta amplitude in a trial with the number of detected theta events for different threshold values (median + x*SD)in both tasks. The high correlation between theta amplitude and number of events at threshold = median +1.5*SD justifies our selection of the same for our analyses.

### Distribution of theta onset times in partial error and full-correct matched trials

#### Flanker Task: Partial Error trials

**Fig. S2.**
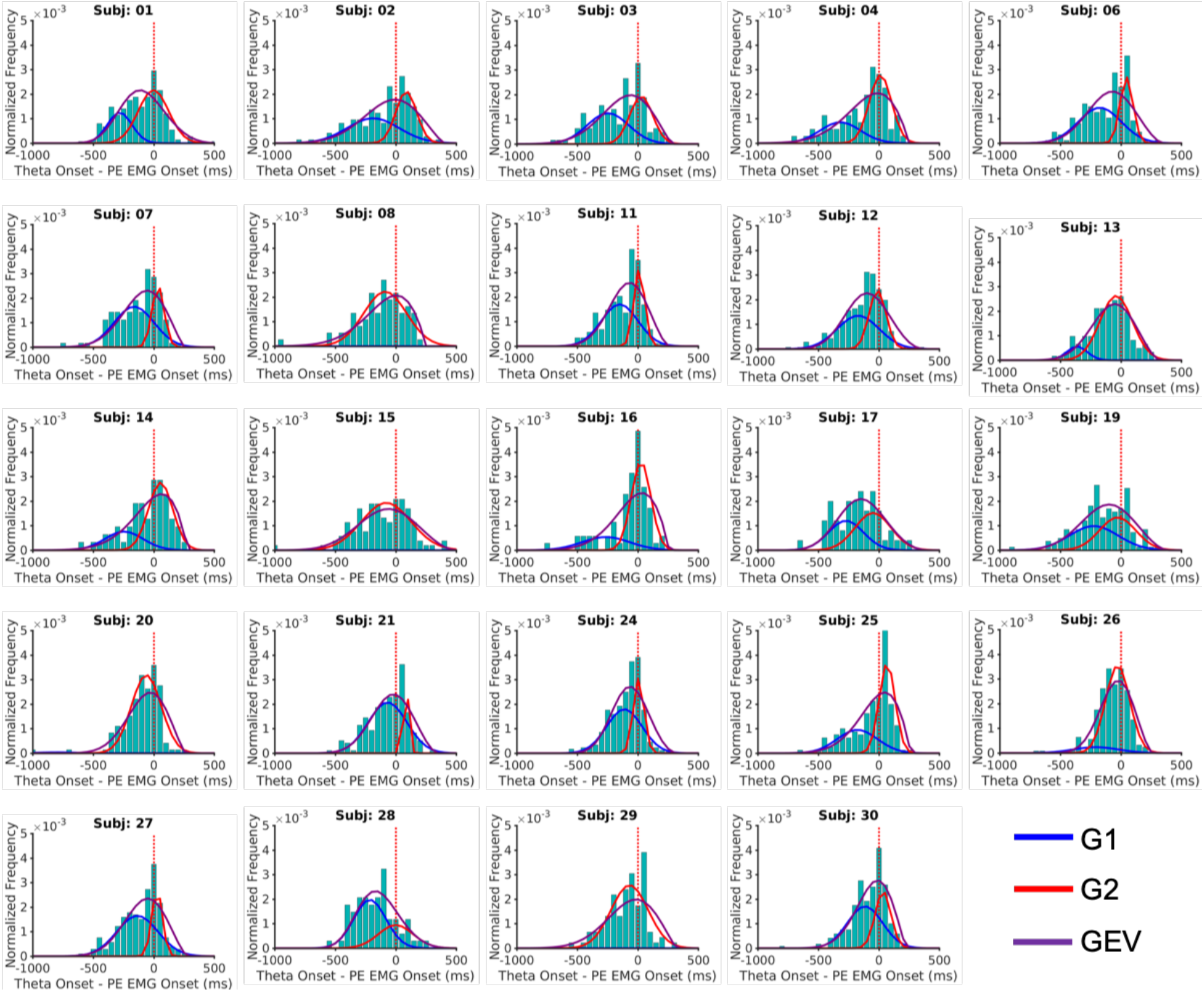
Theta onsets in relation to the partial error EMG onsets in each individual participant in the partial error trials. Each plot has the gaussian mixture model fitted with K = 2 gaussians (G1 and G2) and the fit using the generalized extreme value (GEV) distribution.

#### Flanker Task: Full-correct matched trials

**Fig. S2.**
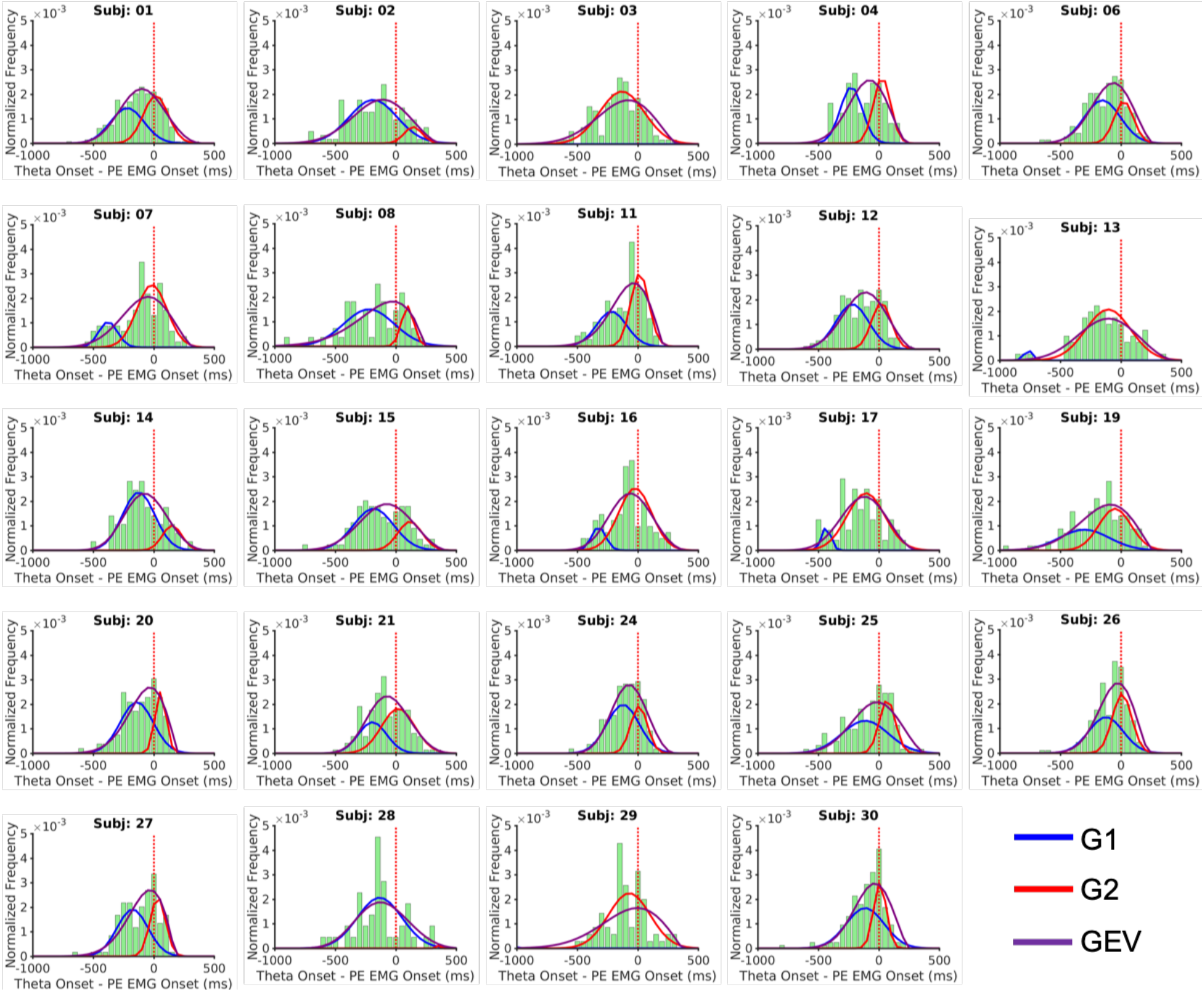
Theta onsets in relation to the surrogate “partial error EMG” onsets in each individual participant in the full-correct matched trials. Each plot has the gaussian mixture model fitted with K = 2 gaussians (G1 and G2) and the fit using the generalized extreme value (GEV) distribution.

#### Simon Task: Partial Error trials

**Fig. S4.**
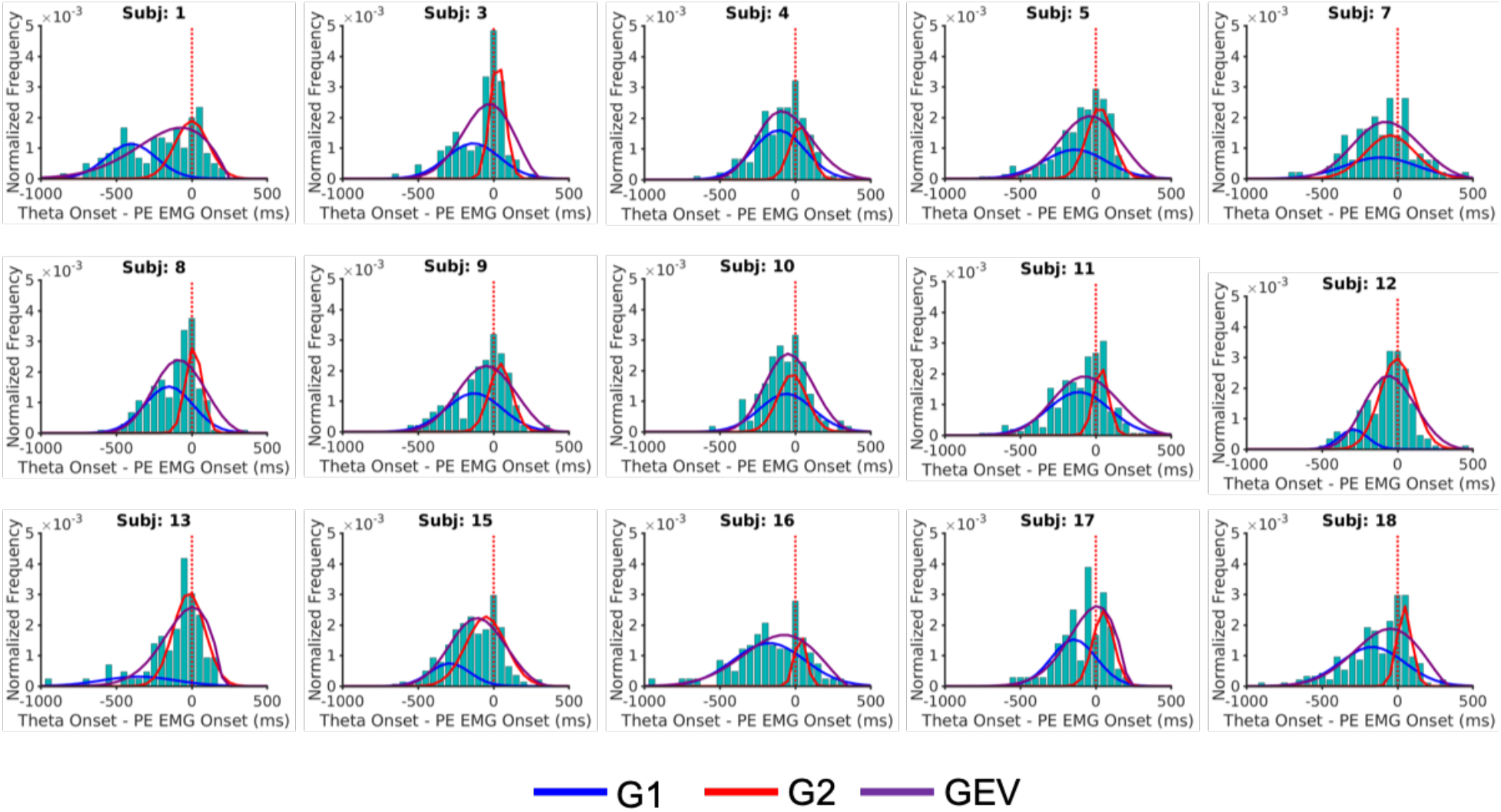
Theta onsets in relation to the partial error EMG onsets in each individual participant in the partial error trials. Each plot has the gaussian mixture model fitted with K = 2 gaussians (G1 and G2) and the fit using the generalized extreme value (GEV) distribution.

#### Simon Task: Full-correct matched trials

**Fig. S5.**
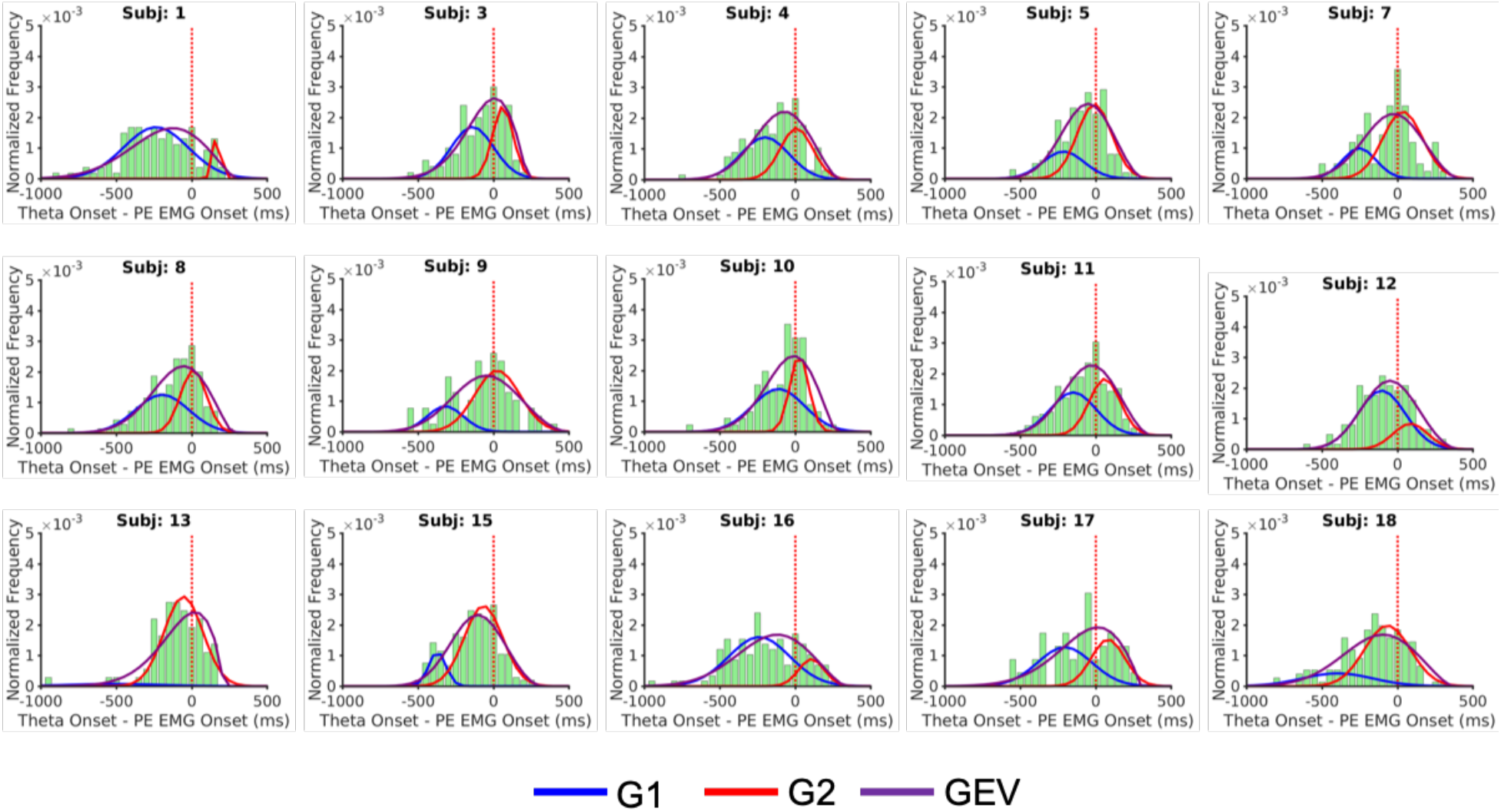
Theta onsets in relation to the surrogate “partial error EMG” onsets in each individual participant in the full-correct matched trials. Each plot has the gaussian mixture model fitted with K = 2 gaussians (G1 and G2) and the fit using the generalized extreme value (GEV) distribution.

### Correlation between theta onsets times (G1 and G2) with EMG timings

**Fig. S6.**
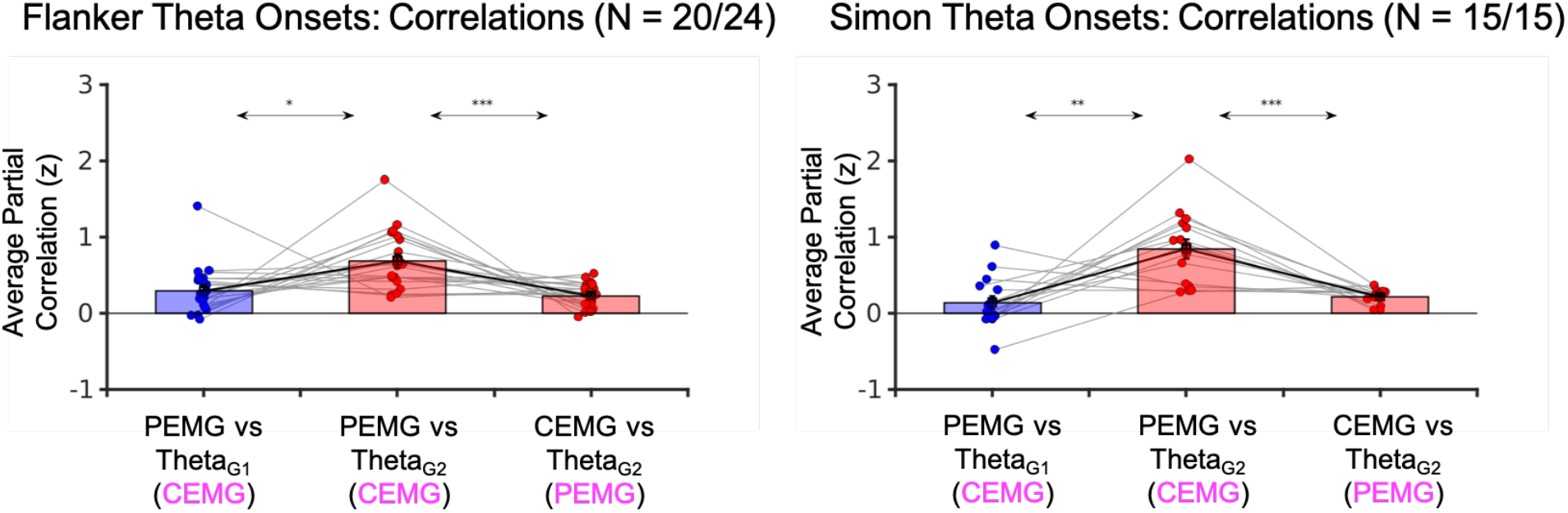
Fisher-transformed partial correlations between the theta onsets belonging to either G1 or G2 and the EMG timings (partial error – PEMG and correct response – CEMG) in the partial error trials for the Flanker and Simon task. The magenta colored variable is the control variable used in the computation of the partial correlation in each case. The correlations are for participants who had more than 5 data points in each of G1 and G2 respectively.

